# GerNA-Bind: Geometric-enhanced RNA-ligand Binding Specificity Prediction with Deep Learning

**DOI:** 10.1101/2025.02.15.638393

**Authors:** Yunpeng Xia, Jiayi Li, Yi-Ting Chu, Jiahua Rao, Jing Chen, Chenqing Hua, Dong-Jun Yu, Xiu-Cai Chen, Shuangjia Zheng

## Abstract

RNA molecules are essential regulators of biological processes and promising therapeutic targets for various diseases. Discovering small molecules that selectively bind to specific RNA conformations remains challenging due to RNA’s structural complexity and the limited availability of high-resolution data. Herein, we introduce GerNA-Bind, a geometric deep learning framework to predict RNA-ligand binding specificity by integrating multi-state RNA-ligand representations and interactions. GerNA-Bind achieves state-of-the-art performance on multiple benchmark datasets and excels in predicting interactions for low-homology RNA-ligand pairs. It achieves a 20.8% improvement in precision for binding site prediction compared to AlphaFold3. Furthermore, it offers informative, well-calibrated predictions with built-in uncertainty quantification. In a large-scale virtual screening application, GerNA-Bind identified 18 structurally diverse compounds targeting the oncogenic MALAT1 RNA, with experimentally confirmed submicromolar affinities. Among them, one lead compound selectively binds the MALAT1 triple-helix, reduces its transcript levels, and inhibits cancer cell migration. These findings highlight GerNA-Bind’s potential as a powerful tool for RNA-focused drug discovery, offering both accuracy and biological insight.

## Introduction

Targeting ribonucleic acids (RNA) with small molecules represents a transformative strategy for modulating biological pathways and developing novel therapeutics. This strategy holds promise for addressing complex diseases by selectively targeting RNA structures^1–3^. Understanding the mechanisms underlying RNA-ligand interactions is critical for advancing RNA-targeted drug discovery and designing effective therapeutics.

Recent efforts have mainly focused on elucidating RNA-ligand binding mechanisms using physics-based approaches. These methods examine forces like electrostatic interactions *π*-*π* stacking interactions, and hydrogen bonding between RNAs and ligands^4–7^. While these methods provide atomistic insights, their predictive power remains bottlenecked by the scarcity of high-resolution structural data, which predominantly relies on resource-intensive techniques like X-ray crystallography or Cryo-EM. Alternative strategies have sought to overcome these challenges using sequence similarity and physicochemical descriptors to predict RNA-ligand binding^8–10^. These methods are simpler and faster but struggle to account for RNA’s structural flexibility. This makes them less effective for predicting binding specificity in dynamic RNA structures. While deep learning models have introduced innovative approaches to predicting RNA-ligand interactions^11–18^, they often face challenges with low-homology RNA-ligand pairs and are less suitable for high-throughput virtual screening due to high model uncertainty. This highlights the need for more robust tools capable of handling diverse RNA structures and interactions effectively.

To address these challenges, we introduce GerNA-Bind, a geometric deep learning framework for predicting RNA-ligand binding. The model combines multiple structural representations with interaction analysis. It uses data from both experimentally resolved structures and high-throughput screening to create functional embeddings. These embeddings capture both graph-based connectivity and three-dimensional spatial interaction relationships. This design improves prediction accuracy and includes uncertainty quantification to assess confidence in the results.

GerNA-Bind demonstrates the transformative potential of geometric deep learning in RNA-ligand interaction research through three key contributions: first, GerNA-Bind achieves state-of-the-art performance on benchmark RNA-ligand interaction datasets, showcasing strong predictive capabilities, especially for low-homology RNA-ligand pairs. Second, by leveraging a triangular geometry module, GerNA-Bind uncovers binding modes across diverse RNA family, providing structural insights into RNA-ligand interactions. Third, application to virtual screening identified 18 structurally diverse binders of the oncogenic MALAT1 lncRNA with submicromolar affinity. One lead compound selectively binds the MALAT1 triple-helix motif, suppresses transcript abundance, and inhibits cancer cell migration. By bridging geometric deep learning with RNA structural biology, GerNA-Bind establishes a robust framework to accelerate the development of RNA-targeted therapeutics. The open release of this tool aims to catalyze innovation in both computational biophysics and translational RNA medicine.

## Results

### Overview of GerNA-Bind

We present the architecture of GerNA-Bind in Fig. 1. GerNA-Bind takes RNA and ligand as inputs, where RNA backbones can be experimentally validated or computationally predicted structures using RNA folding models–RhoFold^19^, and molecular conformations are also derived (Fig. 1A). GerNA-Bind demonstrates high accuracy, robustness and interpretability, making it well-suited for tasks including RNA-ligand binding specificity prediction, RNA-targeting virtual screening, binding site prediction, and binding mechanism analysis (Fig. 1B).

**Figure. 1:**
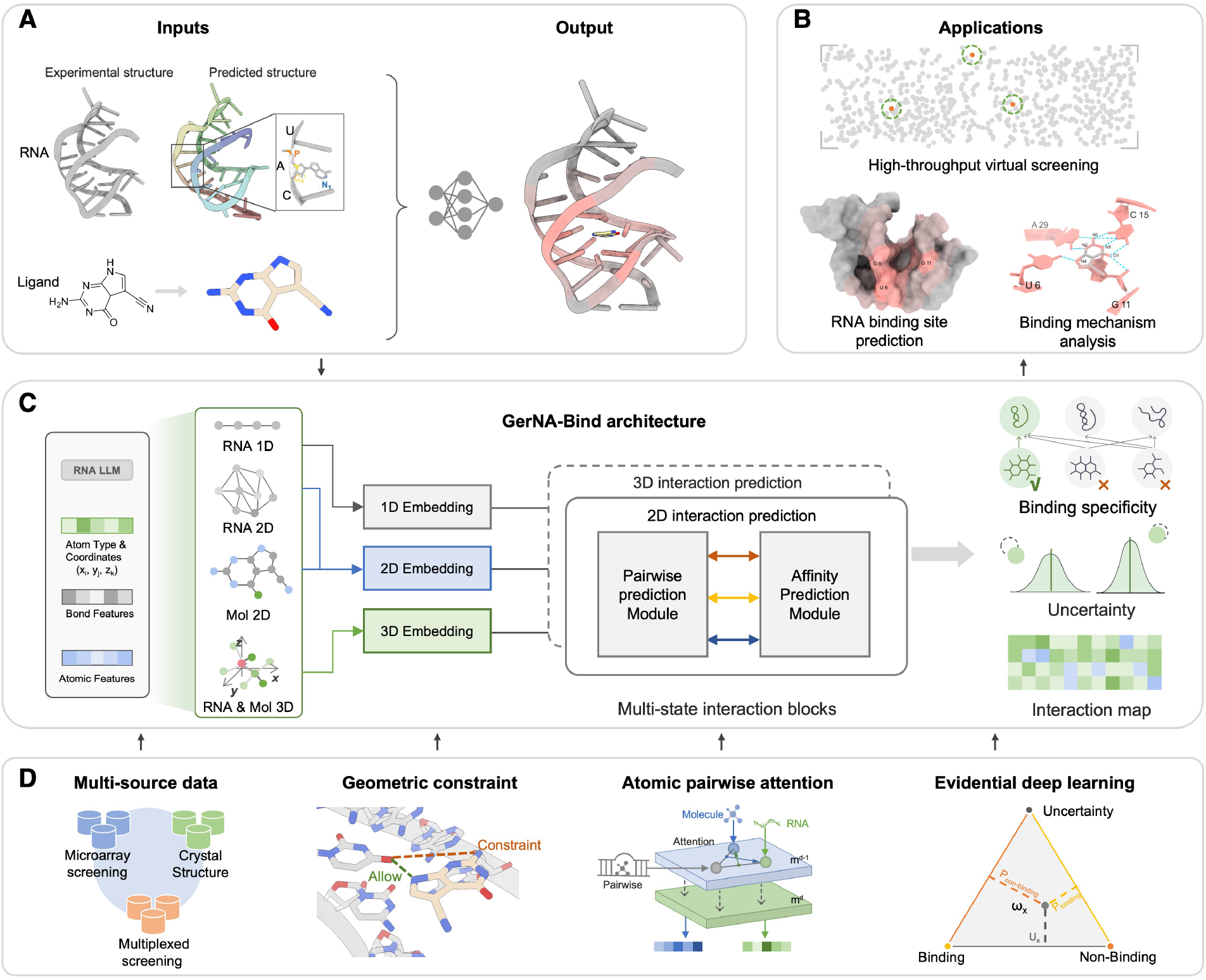
Overview of GerNA-Bind. **(A)** GerNA-Bind takes RNA and ligands as input. RNA is represented by the 3D coordinates of backbone bases and atoms, exemplified by PDB entry 3GCA in the figure. Ligands are represented by their 3D molecular conformations computed using RDKit. **(B)** GerNA-Bind can be applied to various tasks, including RNA-targeting virtual screening, RNA binding site prediction, and binding mechanism analysis. **(C)** GerNA-Bind is a multi-modal neural network for RNA-ligand binding specificity prediction, integrating 1D RNA sequences, 2D RNA secondary structures, 3D RNA conformations, 2D molecular graphs, and 3D molecular conformations. The model captures both 2D graph connectivity and 3D spatial interactions through a pairwise prediction module and an affinity prediction module. It also provides an uncertainty score, offering insights into the confidence of each prediction. **(D)** GerNA-Bind is trained on diverse data sources, including microarray screening, multiplexed screening, and crystal structures. It utilizes geometric constraints to optimize RNA-ligand pairwise interactions and employs an atomic pairwise attention mechanism in the affinity prediction module to identify RNA binding sites. The model integrates evidential deep learning to provide binding specificity predictions along with an uncertainty score to assess prediction confidence.

GerNA-Bind leverages multi-state RNA-ligand representations and interactions by integrating 1D RNA sequences, 2D RNA secondary structures, 3D RNA conformations, 2D molecular graphs, and 3D molecular conformations (Fig. 1C). GerNA-Bind is designed to be simple yet effective, enabling precise RNA-ligand binding specificity predictions through these multi-state inputs. To achieve this, GerNA-Bind employs an equivariant graph transformer to encode 3D RNA conformations, a graph neural network (GNN) to encode 2D RNA secondary structures, and a sequence encoder for 1D RNA sequences. Similarly, graph transformer and GNN encode 3D conformations and 2D graphs for ligands. At its core, GerNA-Bind models both 2D graph connectivity and 3D spatial interactions within RNA-ligand complexes through a dual attention mechanism within an affinity prediction module (Fig. 1D). This module leverages a memory cache to combine pairwise per-base/atom features, RNA representations, and molecular representations, allowing the prediction of affinity, which in turn is used to estimate binding specificity. Additionally, GerNA-Bind incorporates an evidence-parameterized Dirichlet distribution to estimate probabilities and uncertainties for each RNA-ligand complex. This approach enables the model to predict not only binding specificity but also corresponding uncertainty scores, offering valuable insights for decision-making.

To further refine predictions, GerNA-Bind integrates a geometric constraint module, optimizing the base-atom pairwise interaction matrix for RNA and drug-like molecules. This module not only improves pairwise interaction predictions but also enhances interpretability by explicitly predicting the base-atom pairwise interaction matrix. This refinement contributes to a more accurate assessment of RNA-ligand binding specificity derived from the interaction matrix.

### Accurate modeling of RNA-ligand binding specificity

We first evaluated GerNA-Bind’s ability to predict RNA-ligand binding specificity using two public datasets of experimentally validated RNA-ligand interactions: Robin^15^ and Biosensor^20^. To rigorously evaluate model performance, we employed four evaluation splits to simulate in domain and out-of-domain scenarios in which the training and test sets do not share any ligands or RNAs. The detailed description of these two datasets and split strategies can be found in Section Methods.

To benchmark GerNA-Bind’s performance in RNA-ligand binding specificity prediction, we compared it to existing prediction methods, including a state-of-the-art RNA-ligand interaction prediction model (RSAPred^21^), and three adapted protein-ligand interaction prediction models: DeepDTIs^22^, DeepConv-DTI^23^, and GraphDTA^24^. The Area Under the Receiver Operating Characteristic Curve (AUROC) and accuracy, F1 score and Area Under Precision-Recall Curve (AUPRC) were used as the evaluation metric to compare prediction performance across different models.

The results, shown in Fig. 2A, demonstrate that GerNA-Bind significantly outperforms existing models. On the Robin dataset, for the homology and fingerprint-based split, GerNA-Bind achieved a 6.7% improvement in AUROC over the best-performing model, GraphDTA, and a 12.4% improvement over RSAPred. Similar improvements were observed on the Biosensor dataset, where GerNA-Bind outperformed GraphDTA by 9.1% and RSAPred by 12.6%, respectively. Beyond state-of-the-arts performance in the homology and fingerprint-based splits, GerNA-Bind also achieved significant improvements across the other three evaluation splits, demonstrating robust performance and generalizability in RNA-ligand binding specificity prediction. The AUPRC, accuracy, and F1 results are consistent with the AUC, further emphasizing GerNA-Bind’s superior performance.

**Figure. 2:**
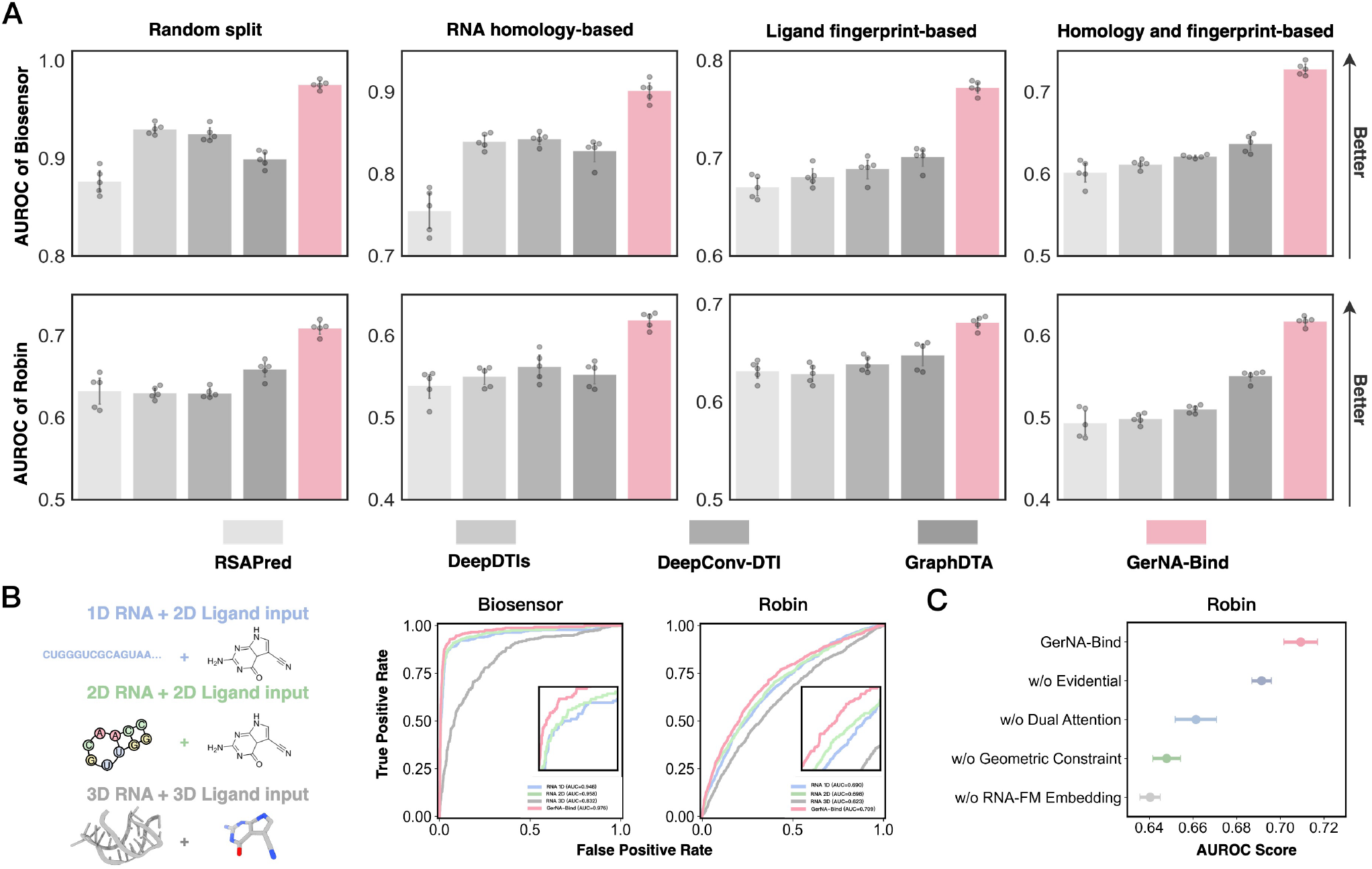
GerNA-Bind in RNA-Ligand Binding Specificity Prediction. **(A)** Comparison of GerNA-Bind to baseline models (RSAPred, DeepDTIs, DeepConv-DTI, and GraphDTA) for RNA-ligand specificity prediction. The AUROC metric is used to evaluate model performance, with GerNA-Bind achieving state-of-the-art results across four different data splits on both the Robin and Biosensor datasets. **(B)** GerNA-Bind leverages a multi-modal framework, modeling interactions between RNA sequences and ligand graphs, RNA graphs and ligand graphs, and RNA structures and ligand structures. Ablation studies highlighting the contribution of multi-state representations in GerNA-Bind. Comparisons to models using single-modality inputs—1D RNA sequences, 2D RNA graphs, or 3D RNA structures—demonstrate that integrating all three RNA modalities significantly enhances GerNA-Bind’s ability to predict RNA-ligand binding specificity. **(C)** Ablation studies of GerNA-Bind without (w/o) corresponding modules in GerNA-Bind with performance measured by AUROC.

The significant advancements achieved by GerNA-Bind can be attributed to its ability to process multi-state RNA-ligand representations and interactions, setting it apart from current methods that primarily focus on single-modality data. As shown in Fig. 2B, GerNA-Bind integrates multi-state RNA and molecular data, combining sequence, graph, and structural information for enhanced predictive accuracy. To validate the benefits of multi-state representations, we conducted ablation studies on the random split of both the Biosensor and Robin datasets, as illustrated in Fig. 2C. When compared to ablation models that employ only single-modality data—such as sequence, graph, or structure representation—GerNA-Bind achieved significant performance gains, with AUROC scores increasing by 14% on the Biosensor dataset and 9.7% on the Robin dataset. These results underscore that incorporating multi-state RNA-ligand representations, including sequences, graphs, and structures, establishes GerNA-Bind as a highly effective framework for RNA-ligand binding specificity prediction.

### Uncertainty quantification of RNA-ligand interaction prediction

GerNA-Bind integrates uncertainty estimation as a performance confidence measure to support decision-making. This capability enhances virtual screening processes for identifying promising drug candidates. GerNA-Bind achieves this by leveraging an evidence-based deep learning framework, which assigns an uncertainty score to each RNA-ligand binding prediction. Details of the uncertainty estimation module are discussed in Section Methods.

To evaluate the reliability of binding predictions and the utility of uncertainty estimations for decision-making, we first quantitatively analyzed the relationship between uncertainty estimates and prediction performance (AUROC scores) using the Robin dataset (Fig. 3). Here, predictions were ranked in ascending order of uncertainty and grouped into percentiles, with cumulative AUROC calculated for each group. In Fig. 3A, GerNA-Bind showed a strong correlation between uncertainty estimates and prediction performance, outperforming ensemble and dropout approaches. In a random data split analysis (Fig. 3B), GerNA-Bind achieved a Spearman correlation of 0.96 between uncertainty estimates and prediction performance, compared to 0.79 for ensemble learning and 0.82 for dropout. These results underscore the robustness of GerNA-Bind’s predictions and uncertainty estimates, demonstrating its potential to guide virtual screening and experimental validation in RNA-targeting drug discovery.

**Figure. 3:**
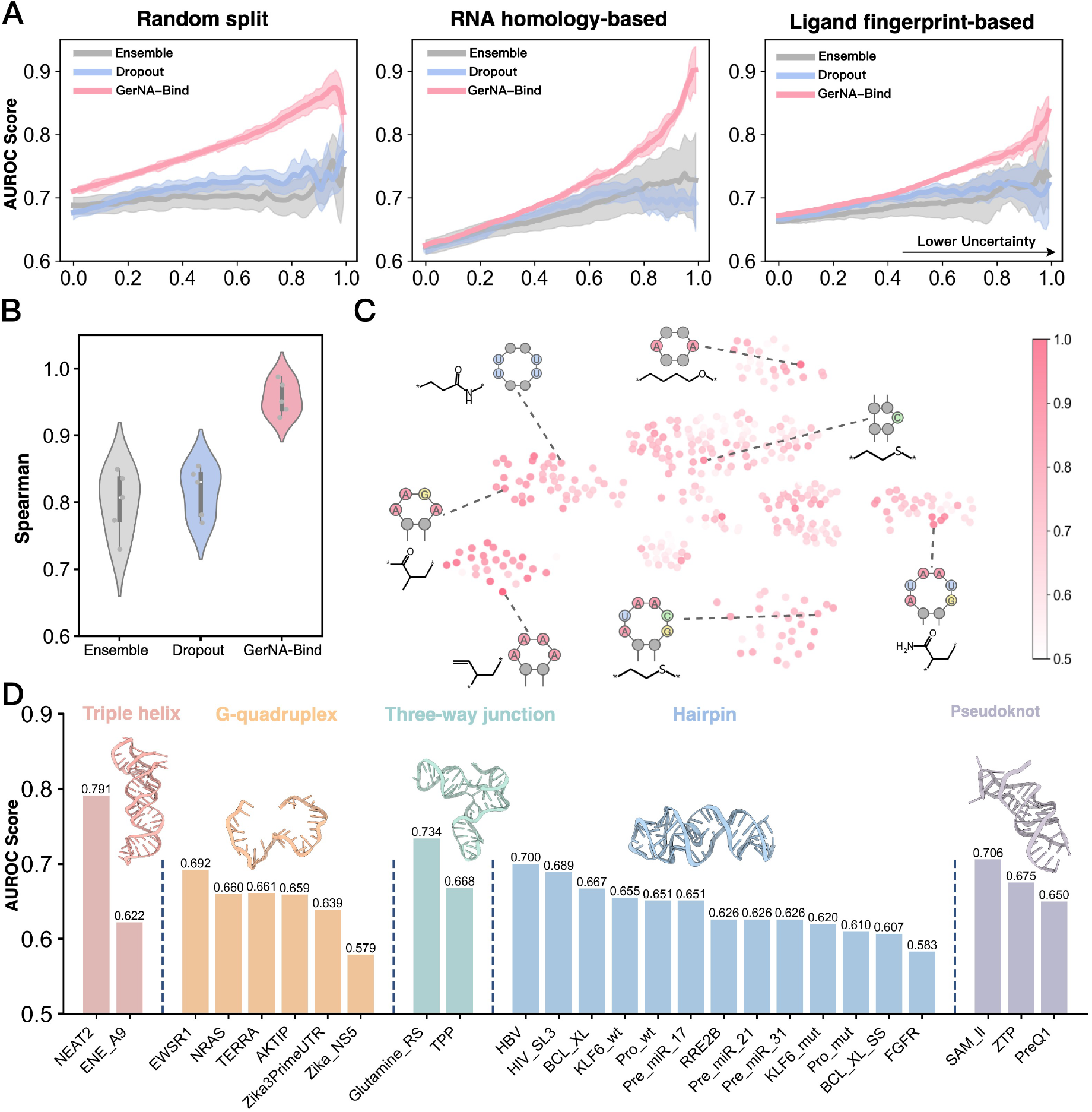
Uncertainty quantification of RNA-ligand interaction using GerNA-Bind. **(A)** Correlation between uncertainty estimates and binding prediction performance. GerNA-Bind is compared to ensemble learning and the dropout approach across random, homology-based, and fingerprint-based splits. GerNA-Bind demonstrates robust predictions, with high AUROC scores associated with low uncertainty. **(B)** Spearman correlation analysis in the random split setting, revealing a strong positive correlation between model confidence and binding prediction performance in GerNA-Bind. **(C)** Classification performance across sub-test sets in the Robin dataset. Each sub-test set represents RNA-ligand complexes defined by RNA secondary structures and drug-like molecule substructures. Darker color intensities, indicating higher AUROC values, are correlated with improved performance. **(D)** GerNA-Bind’s performance across different RNA structural types, including triple helices, G-quadruplexes, three-way junctions, hairpins, and pseudoknots. The figure shows AUROC scores for binding specificity prediction across 26 RNA targets.

In addition to correlation analysis, we evaluated GerNA-Bind’s ability to predict RNA pocket–small molecule substructure compatibility across diverse RNA architectures using the Robin dataset, focusing on structural matches between defined RNA binding pockets and ligand pharmacophore motifs. First, we refined the dataset by extracting 27 common substructures from drug-like molecules that appeared in more than 1,000 of the 24,572 molecules in the Robin dataset. We then used Inforna^8^ to identify 19 biologically relevant RNA secondary structure motifs, resulting in a subset of 513 combinations for focused assessment of binding prediction performance. Sub-test results (Fig. 3C) revealed that darker color intensities, indicating higher AUROC, were associated with improved prediction performance for biologically significant RNA motifs. Notably, adenine- and uracil-enriched regions exhibit a propensity to interact with small molecules bearing amine (-NH2) or hydroxyl (-OH) groups, indicating the potential for hydrogen bonding interactions that may contribute to the stabilization of binding and further enhance binding affinity. These findings highlight the ability of GerNA-Bind to identify RNA motifs that have the potential to interact with specific small molecules, providing valuable insights into RNA-ligand binding. Additionally, we evaluated GerNA-Bind’s performance across various RNA structural types, including triple helices, G-quadruplexes, three-way junctions, hairpins, and pseudoknots (Fig. 3D). GerNA-Bind achieved its best performance on NEAT2 RNA, with an AUROC of 0.791, and demonstrated consistent performance (AUROC around 0.6) across other RNA types. These results highlight GerNA-Bind’s stability and robustness in RNA-ligand binding prediction across various RNA structures, emphasizing its practical usage for drug discovery.

GerNA-Bind’s evidence-based learning framework help identify active bases and atoms critical for RNA-ligand binding, delivering interpretable predictions and improving generalizability to unseen RNA-ligand interactions. Unlike traditional alignment- or similarity-based methods that may struggle with novel complexes, GerNA-Bind excels in identifying binding specificities in unexplored chemical and biological spaces. This positions GerNA-Bind as a powerful tool for advancing virtual screening and decision-making in RNA-targeting drug discovery.

### GerNA-Bind’s superiority in RNA binding site identification

GerNA-Bind not only predicts RNA-ligand binding specificity but also directly models their interactions. By leveraging its geometric constraint module, GerNA-Bind learns a pairwise contact matrix that captures RNA-ligand interactions (Fig. 1D). To validate its capability, GerNA-Bind was fine-tuned on the Hariboss dataset^25^, comprising 359 experimentally validated RNA-ligand crystal structures with accurate interaction data. The model was pre-trained on the Robin dataset for 30 epochs and then fine-tuned on the Hariboss dataset, using only RNA-ligand complexes from 2020 and earlier for training (318 complexes), with the remaining 19 and 22 complexes from 2021 and later used for validation and testing, respectively. In this setting, each entry in the RNA-ligand contact matrix represents the minimal atomic distance between each RNA base and each atom of the drug-like molecule. The pairwise contact matrix was also used to estimate the binding affinity variance between RNA targets and drug-like molecules, improving prediction performance.

Our model was benchmarked against leading structural prediction frameworks, including RNASite^26^, Chai-1^27^, and Alphafold3^18^, using comprehensive metrics such as Recall, Precision, F1-score, AUPRC, and MCC (Fig. 4A). On these evaluations, our model outperformed the competition, delivering a 20.8% increase in precision, a 13% improvement in F1-score, a 10.6% boost in AUPRC, and a 15.6% enhancement in MCC compared to Alphafold3, showcasing its exceptional accuracy in predicting RNA-binding sites. Moreover, to evaluate its performance across diverse RNA structural categories, we analyzed the 22 RNA structures in the test set. Impressively, our model achieved the highest AUROC scores in four out of five RNA structural classes (Fig. 4B), underscoring its broad adaptability and precision. Binding site predictions were visualized by extracting maximum values from the predicted interaction matrix for each RNA base, representing the likelihood of it being a binding site, and these were rendered using ChimeraX. Five RNA-ligand complexes from distinct structural clusters further demonstrated the model’s capacity to accurately predict targets across varied RNA configurations (Fig. 4C). Notably, the predicted binding sites formed effective hydrogen bonds with small molecules, a predominant interaction type in RNA-ligand interactions (Fig. 4D). Collectively, these findings establish GerNA-Bind as a cutting-edge tool for pinpointing RNA-ligand binding sites with unmatched precision.

**Figure. 4:**
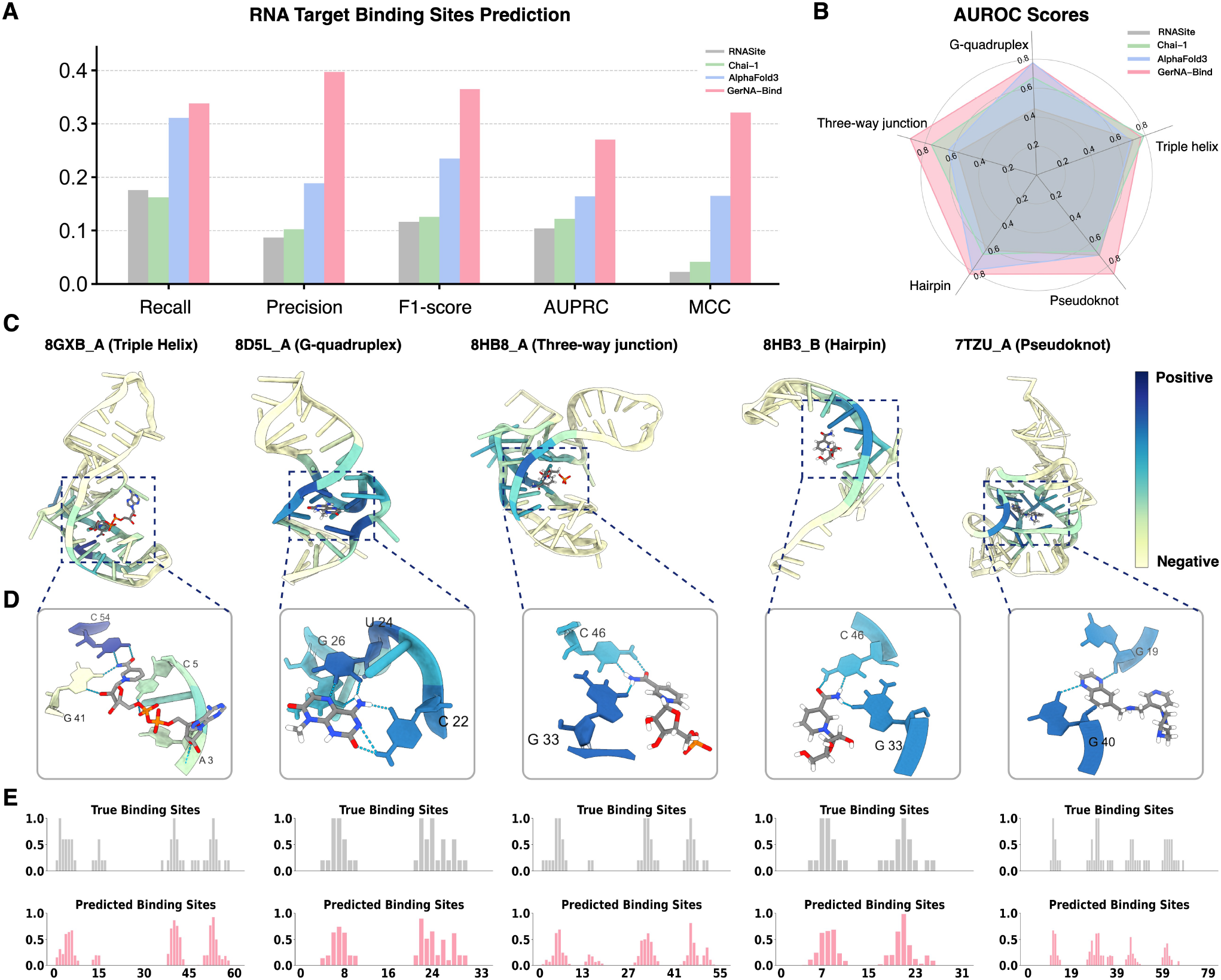
GerNA-Bind Validation for RNA Binding Site Identification. **(A)** GerNA-Bind effectively predicts RNA target binding sites, with its performance significantly surpassing that of models such as RNASite, Chai-1, and AlphaFold3. **(B)** The AUROC scores illustrate the model’s ability to predict binding sites across five distinct RNA structures: Triple Helix, G-Quadruplex, Three-Way Junction, Hairpin, and Pseudoknot. **(C)** Specific examples corresponding to these five RNA structures are provided, with darker regions denoting a higher likelihood of a base being classified as a binding site by the model. **(D)** Through the model’s analysis of the local structures in the above cases, we demonstrated that the predicted RNA binding sites establish hydrogen bond interactions with small molecules.

### Virtual screening of MALAT1 triple helix with GerNA-Bind

A critical challenge in RNA-targeting drug discovery is to come up with small molecules that have good efficacy while being selective. We reasoned that GerNA-Bind could enable assessment of the accuracy and pre-organization of designed active sites by generating ensembles of models of small molecules and interacting sidechain placements much more rapidly than physics-based methods and without the limitations of requiring experimental structural data.

To demonstrate the effectiveness of our newly developed method, we conducted a large-scale virtual screening campaign against a therapeutic-related RNA target, oncogenic MALAT1 (metastasis-associated lung adenocarcinoma transcript 1) lncRNA. MALAT1 plays a role in metastasis and tumor growth in many types of cancer^28,29^. Depleting MALAT1 through different approaches shows strong antiproliferative effects. For this reason, MALAT1 is seen as a potential target for cancer treatment^30,31^. One major challenge in developing MALAT1-targeting drugs is finding molecules that selectively bind to the MALAT1 triple helix. This structure helps accumulation of the transcript, and high levels of MALAT1 are linked to several cancers^32^. We aimed to find structurally diverse compounds that could selectively bind to this triple helix. So far, only one study has reported a molecule with a diphenylfuran structure that can act on it^33^.

We began by using a substructure matching algorithm based on common molecular substructures identified in the RNA-targeting small molecule dataset. This dataset includes active small molecules from Robin, Biosensor, and Hariboss datasets. Using this approach, we excluded the majority of molecules that lacked relevant substructures from the Topscience refine set, a commercial compound library containing 8,357,246 compounds. As a result, we created an RNA-targeting focused library consisting of 21,659 compounds. Next, we applied predictive ensembles of GerNA-Bind to identify triple helix-binding molecules (score > 0.8 and uncertainty < 0.15) and stem-loop non-binding molecules (score < 0.2 and uncertainty < 0.15). We applied a similarity filter against active molecules in the RNA targeting small molecule dataset using a tanimoto similarity threshold of < 0.6 and excluded similar RNA sequences (80% homology) from the GerNA-Bind training dataset to prevent potential data leakage. This process narrowed the collection down to 140 and 49 compounds, respectively. Following this, we performed clustering based on molecular similarity. From the resulting clusters, we selected the highest-ranking molecules that showed favorable interactions and geometry. This process ultimately led to the identification of 28 promising candidate compounds (Fig. 5A).

**Figure. 5:**
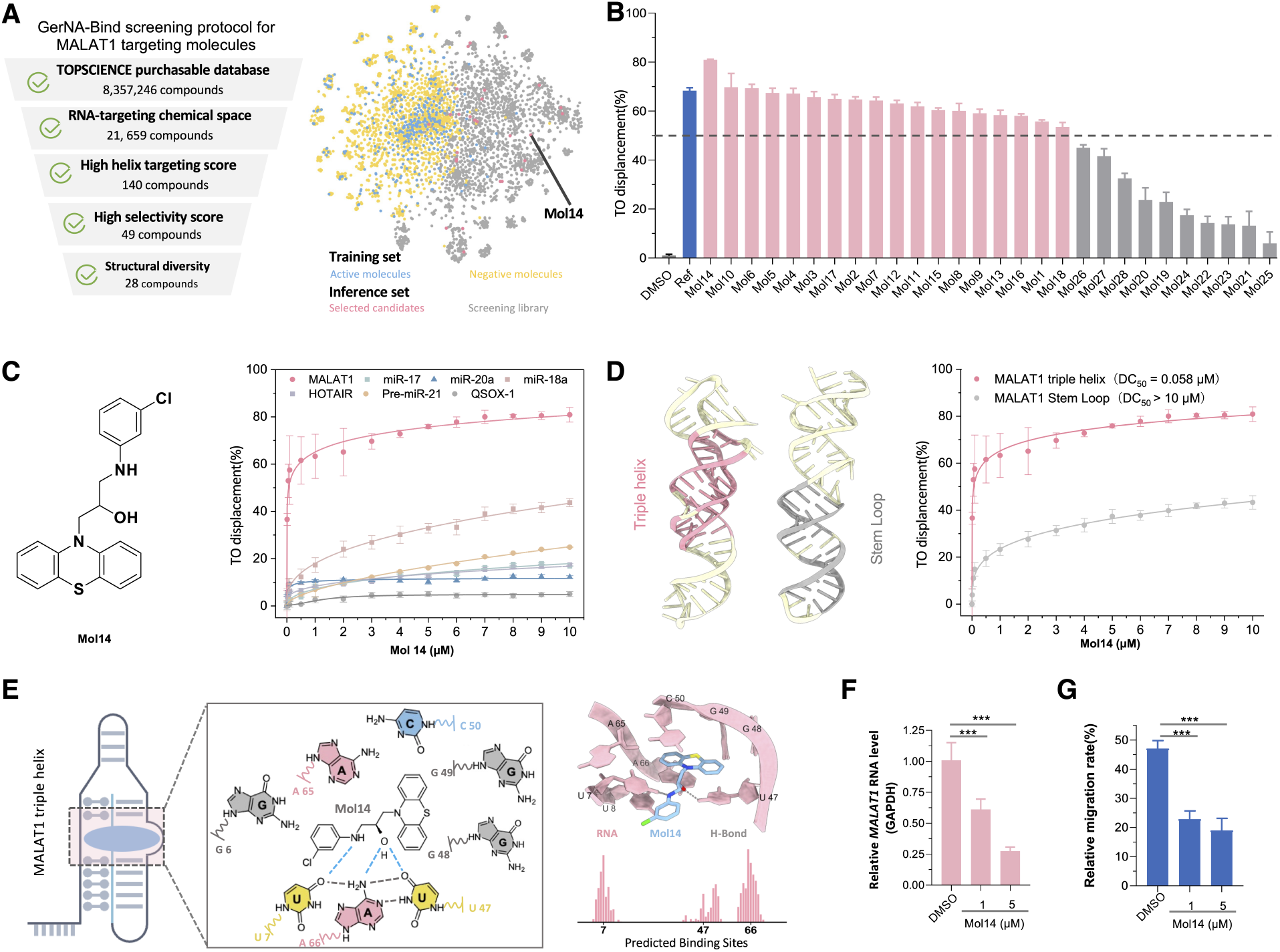
GerNA-Bind assists wet-lab experiments for RNA-targeting drug discovery. **(A)** Left: The detailed process by which the model screens candidate drugs for MALAT1. Right: A schematic representation of the spatial distribution of the drug molecules; **(B)** The screening results of 28 GerNA-Bind identified molecules using TO displacement assays for MALAT1 at a concentration of 10 µM; **(C)** Dose–response curve of TO displacement from various RNAs upon the addition of increasing concentrations of Mol14; **(D)** Dose–response curve of TO displacement from MALAT1 triple helix or stem loop upon the addition of increasing concentrations of Mol14; **(E)** Analysis of the interaction between MALAT1 and the small molecule Mol14. Left: The RNA three-helix bundle structure. Center: A close-up view of the RNA secondary structure within the binding pocket. Top-right: AlphaFold3-predicted structure with RNA in red, Mol14 in blue, and hydrogen bonds in gray. Bottom-right: Binding site predictions from the GerNA-Bind model; **(F)** MALAT1 RNA levels in A549 cells treated with Mol14; **(G)** Evaluation of cell migration using the wound healing assay. The data are presented as mean ± SD from three independent experiments, and statistical significance is determined by the t-test with the following annotations: (ns) not significant, (*) P < 0.05, (**) P < 0.01, and (***) P < 0.001.

To evaluate the binding affinity of the 28 candidates for the oncogenic MALAT1 RNA, thiazole orange (TO) displacement assays were performed. Compound 16 (Ref), a reported MALAT1 ligand, was used as a positive control^34^. In the TO displacement assay, TO fluoresced upon binding to RNA, and the fluorescence intensity decreased as compounds were added to compete for the RNA binding site^35–38^. A higher displacement ratio indicated a stronger affinity of the compounds for RNA. Among these compounds, 64% (Mol1–18) displaced more than 50% of TO from MALAT1 RNA, indicating a strong binding affinity for MALAT1 RNA (pink bars; Fig. 5B). Notably, three novel molecules Mol14, Mol10, and Mol6 demonstrated better displacement ratio (with values of 80.9%, 69.8% and 69.3%, respectively) than ref (68.4%). Based on the above data, the most potent compound, Mol14 (a phenothiazine derivative), was selected for further detailed investigations.

Several disease-related RNAs, including those implicated in cancer, were selected to investigate the selectivity of Mol14 towards MALAT1. These RNAs included miR-17, miR-20a, miR-18a, HOTAIR, Pre-miR-21, and QSOX-1^39–42^. The binding affinity was reported as the competitive displacement value (DC50), which represents the concentration of small molecules required to achieve 50% displacement of TO^35,36^. As depicted in Fig. 5C, Mol14 exhibited high specificity for MALAT1 (DC50 = 58 nM) and weak binding to the other six RNAs (DC50 > 10 µM).

Given the exciting potential of Mol14 as a selective ligand for MALAT1 RNA, we further investigated the binding mode of Mol14 with MALAT1 RNA. The bases forming the triple helix were mutated to construct non-triplex-forming stem loop, and then TO displacement assay was performed to compare the binding affinity of Mol14 to these two RNA structures. It was found that Mol14 exhibited extremely weak binding with MALAT1 stem loop, suggesting that Mol14 bound to MALAT1 RNA at the triple helix site (Fig. 5D). The protruding guanines at positions 48 and 49, along with the surrounding triple-helix architecture, form a unique binding pocket (Fig. 5E). Notably, Mol14 forms hydrogen bonds with U7, U47, and A66—-nucleotides that directly contribute to the triple-helix structure in MALAT1 (predicted by AlphaFold3). Moreover, GerNA-Bind accurately predicts U7 and A66 as key binding sites. These findings not only explain the significant loss of binding affinity observed when mutations disrupt the triple-helix at these positions, but also provide insight into Mol14’s specific interaction with the MALAT1 triple-helix region, offering a structural rationale for its exceptional target specificity.

The above results demonstrated the selective binding of Mol14 to MALAT1. Considering that small molecule ligands targeting MALAT1 can disrupt the RNA structure and then cause its reduction in cells^33,34^, we investigated the impact of Mol14 on the levels of MALAT1 in A549 lung adenocarcinoma cells by qPCR. As shown in Fig. 5F, Mol14 decreased the MALAT1 mRNA levels in a dose-dependent manner. In addition, we found that the cell migration rate was significantly reduced with the treatment of Mol14 in Fig. 5G. As MALAT1 is important for tumor cell motility and its loss will affect the migration, this result also revealed that Mol14 changed the RNA levels. Collectively, these results proved that Mol14 bound MALAT1 effectively and specifically. Above findings highlight the potential of GerNA-Bind for RNA-targeting drug discovery.

## Discussion

GerNA-Bind is designed to model RNA-ligand interactions and predict RNA-ligand binding specificity, with the aim of accelerating RNA-targeting virtual screening and drug discovery. Traditional approaches heavily rely on high-throughput screening and high-resolution structural data, which are expensive and limit generalizability across diverse RNA types. Recent deep learning methods have sought to improve generalizability, but while they show better performance in binding prediction, they often fail to model RNA-ligand interactions explicitly, making their predictions less interpretable. GerNA-Bind addresses these limitations by not only improving binding prediction performance but also enhancing generalizability and interpretability through the modeling of RNA-ligand interactions at the granularity of per-base and per-atom pairwise contact matrices.

GerNA-Bind is a geometric deep learning framework that integrates multi-state RNA-ligand representations, including 1D sequences, 2D graphs, and 3D structures. Additionally, it incorporates an evidence-based learning and uncertainty estimation module, which enhances the accuracy, interpretability, and reliability of its predictions. Computationally, GerNA-Bind achieves state-of-the-art performance in RNA-ligand binding specificity prediction on public datasets and extends its capabilities to accurately predict RNA-ligand binding sites. Experimentally, in wet-lab validation, GerNA-Bind identified 18 diverse hits and one lead molecule that selectively target the oncogenic MALAT1 triple helix with functional validation. This highlights GerNA-Bind’s potential to accelerate virtual screening and facilitate RNA-targeting drug discovery.

While GerNA-Bind demonstrates state-of-the-art performance in RNA-ligand binding specificity prediction, it has certain limitations. The model is specialized for RNA-ligand interactions and does not currently support the modeling of more complex interactions, such as RNA-protein^17,43^ or polymer-polymer interactions, or chemically modified RNA bases. Additionally, its performance can be constrained by the availability of high-throughput screening RNA-ligand data. Integrating more experimental data from structural analysis tools like PLIP^44^ or leveraging transfer learning from protein-small molecule interaction datasets could potentially address this limitation. Furthermore, while GerNA-Bind provides robust predictions, its utility is enhanced when integrated with generative models like RFDiffusionAA^43^ or AlphaFold3^18^, which require quality filtering and specificity scoring for generated complexes. Future developments could focus on expanding its applicability to other macromolecular interactions, incorporating additional experimental and computational datasets, and optimizing its integration with generative frameworks to further advance RNA-targeting drug discovery.

In summary, GerNA-Bind represents a robust computational framework for RNA-ligand binding specificity prediction, addressing important challenges in virtual screening and drug selection for RNA-targeting therapeutics. Its multi-modal architecture effectively combines experimentally validated and computationally predicted RNA structures, offering both predictive accuracy, interpretability, and generalizability. GerNA-Bind’s integration with future generative modeling approaches and advancements in multi-macromolecular interaction modeling holds the potential to significantly advance RNA-targeting drug discovery. As the field progresses, GerNA-Bind can set a foundation for computationally driven structural biology, offering new avenues for therapeutic interventions targeting RNA in disease contexts.

## Data Availability

We utilized publicly accessible datasets as detailed in the Methods section. The preprocessed Robin and Biosensor datasets are available at Zenodo (https://zenodo.org/records/14808549). RNA structural features were derived using three established tools: RNA-FM (https://github.com/ml4bio/RNA-FM) for one-dimensional sequence embeddings, RNAfold (http://rna.tbi.univie.ac.at) for secondary structure predictions, and RhoFold (https://github.com/ml4bio/RhoFold) for three-dimensional structural modeling. Structural data for fine-tuning the model’s RNA-ligand interaction prediction capability were obtained from Hariboss (https://hariboss.pasteur.cloud/), with corresponding RNA tertiary structures retrieved from the RCSB Protein Data Bank (https://www.rcsb.org/).

## Code availability

The source code and the pretrained model weights of GerNA-Bind is freely available at https://github.com/GENTEL-lab/GerNA-Bind.

## Acknowledgements

This study has been supported by the National Natural Science Foundation of China [62402314, 62372234, 62072243, 22207135], the Guangdong Basic and Applied Basic Research Foundation [2023A1515012616], the Young Elite Scientists Sponsorship Program by CAST [2023QNRC001], the Science and Technology Commission of Shanghai Municipality [24510714300], and the project from Smart Medical Innovation Technology Center-GDUT [ZYZX24-011]. S. Z. acknowledges funding from the Asian Young Scientist Fellowship.

## Author contributions statement

S.Z. conceived and supervised the project. Y.X., J.R. and S.Z. contributed to the algorithm implementation. J.L. and Y.X. performed the data preprocessing. Y.X., S.Z., J.L. and D.Y. contributed to the visualization implementation. Y.C., J.C. and X.C. conducted the wet-lab experiments. S.Z.,Y.X., X.C., C.H. and J.L. wrote the manuscript. All authors were involved in the discussion and proofreading.

## References

1. Warner, K. D., Hajdin, C. E. & Weeks, K. M. Principles for targeting rna with drug-like small molecules. Nat. reviews Drug discovery 17, 547–558 (2018).

2. Childs-Disney, J. L. et al. Targeting rna structures with small molecules. Nat. Rev. Drug Discov. 21, 736–762 (2022).

3. Hopkins, A. L. & Groom, C. R. The druggable genome. Nat. reviews Drug discovery 1, 727–730 (2002).

4. Corley, M., Burns, M. C. & Yeo, G. W. How rna-binding proteins interact with rna: molecules and mechanisms. Mol. cell 78, 9–29 (2020).

5. Xie, Y.-c., Eriksson, L. A. & Zhang, R.-b. Molecular dynamics study of the recognition of atp by nucleic acid aptamers. Nucleic acids research 48, 6471–6480 (2020).

6. Tong, Y. et al. Programming inactive rna-binding small molecules into bioactive degraders. Nature 618, 169–179 (2023).

7. Koehn, J. T., Felder, S. & Weeks, K. M. Innovations in targeting rna by fragment-based ligand discovery. Curr. opinion structural biology 79, 102550 (2023).

8. Disney, M. D. et al. Inforna 2.0: a platform for the sequence-based design of small molecules targeting structured rnas. ACS chemical biology 11, 1720–1728 (2016).

9. Sun, S., Yang, J. & Zhang, Z. Rnaligands: a database and web server for rna–ligand interactions. Rna 28, 115–122 (2022).

10. Donlic, A. et al. R-bind 2.0: an updated database of bioactive rna-targeting small molecules and associated rna secondary structures. ACS chemical biology 17, 1556–1566 (2022).

11. Cai, Z., Zafferani, M., Akande, O. M. & Hargrove, A. E. Quantitative structure–activity relationship (qsar) study predicts small-molecule binding to rna structure. J. medicinal chemistry 65, 7262–7277 (2022).

12. Rekand, I. H. & Brenk, R. Drugpred_rna—a tool for structure-based druggability predictions for rna binding sites. J. chemical information modeling 61, 4068–4081 (2021).

13. Grimberg, H. et al. Machine learning approaches to optimize small-molecule inhibitors for rna targeting. J. Cheminformatics 14, 4 (2022).

14. Deng, Z. et al. Predicting ligand-rna binding using e3-equivariant network and pretraining. In Machine learning for structural biology workshop, NeurIPS (2022).

15. Yazdani, K. et al. Machine learning informs rna-binding chemical space. Angewandte Chemie 135, e202211358 (2023).

16. Sun, S. & Gao, L. Contrastive pre-training and 3d convolution neural network for rna and small molecule binding affinity prediction. Bioinformatics btae155 (2024).

17. Baek, M. et al. Accurate prediction of protein–nucleic acid complexes using rosettafoldna. Nat. methods 21, 117–121 (2024).

18. Abramson, J. et al. Accurate structure prediction of biomolecular interactions with alphafold 3. Nature 1–3 (2024).

19. Shen, T. et al. E2efold-3d: end-to-end deep learning method for accurate de novo rna 3d structure prediction. arXiv preprint arXiv:2207.01586 (2022).

20. Townshend, B., Kaplan, M. & Smolke, C. D. Highly multiplexed selection of rna aptamers against a small molecule library. Plos one 17, e0273381 (2022).

21. Krishnan, S. R., Roy, A. & Gromiha, M. M. Reliable method for predicting the binding affinity of rna-small molecule interactions using machine learning. Briefings Bioinforma. 25, bbae002 (2024).

22. Wen, M. et al. Deep-learning-based drug–target interaction prediction. J. proteome research 16, 1401–1409 (2017).

23. Lee, I., Keum, J. & Nam, H. Deepconv-dti: Prediction of drug-target interactions via deep learning with convolution on protein sequences. PLoS computational biology 15, e1007129 (2019).

24. Nguyen, T. et al. Graphdta: predicting drug–target binding affinity with graph neural networks. Bioinformatics 37, 1140–1147 (2021).

25. Panei, F. P., Torchet, R., Menager, H., Gkeka, P. & Bonomi, M. Hariboss: a curated database of rna-small molecules structures to aid rational drug design. Bioinformatics 38, 4185–4193 (2022).

26. Su, H., Peng, Z. & Yang, J. Recognition of small molecule–rna binding sites using rna sequence and structure. Bioinformatics 37, 36–42 (2021).

27. Discovery, C. et al. Chai-1: Decoding the molecular interactions of life. bioRxiv 10.1101/2024.10.10.615955 (2024). https://www.biorxiv.org/content/early/2024/10/15/2024.10.10.615955.full.pdf.

28. Gutschner, T. et al. The noncoding rna malat1 is a critical regulator of the metastasis phenotype of lung cancer cells. Cancer research 73, 1180–1189 (2013).

29. Ji, P. et al. Malat-1, a novel noncoding rna, and thymosin β 4 predict metastasis and survival in early-stage non-small cell lung cancer. Oncogene 22, 8031–8041 (2003).

30. Guo, F. et al. Inhibition of metastasis-associated lung adenocarcinoma transcript 1 in caski human cervical cancer cells suppresses cell proliferation and invasion. Acta Biochim Biophys Sin 42, 224–229 (2010).

31. Tano, K. et al. Malat-1 enhances cell motility of lung adenocarcinoma cells by influencing the expression of motility-related genes. FEBS letters 584, 4575–4580 (2010).

32. Arun, G. et al. Differentiation of mammary tumors and reduction in metastasis upon malat1 lncrna loss. Genes & development 30, 34–51 (2016).

33. Donlic, A. et al. Discovery of small molecule ligands for malat1 by tuning an rna-binding scaffold. Angewandte Chemie 130, 13426–13431 (2018).

34. Abulwerdi, F. A. et al. Selective small-molecule targeting of a triple helix encoded by the long noncoding rna, malat1. ACS chemical biology 14, 223–235 (2019).

35. Rocca, R. et al. Hit identification of novel small molecules interfering with malat1 triplex by a structure-based virtual screening. Arch. der Pharmazie 356, 2300134 (2023).

36. Krishnamurthy, M., Schirle, N. T. & Beal, P. A. Screening helix-threading peptides for rna binding using a thiazole orange displacement assay. Bioorganic & medicinal chemistry 16, 8914–8921 (2008).

37. Wang, Z.-F. et al. The hairpin form of r (g4c2) exp in c9als/ftd is repeat-associated non-atg translated and a target for bioactive small molecules. Cell chemical biology 26, 179–190 (2019).

38. Tran, T. & Disney, M. D. Identifying the preferred rna motifs and chemotypes that interact by probing millions of combinations. Nat. communications 3, 1125 (2012).

39. Liu, X. et al. Targeted degradation of the oncogenic microrna 17-92 cluster by structure-targeting ligands. J. Am. Chem. Soc. 142, 6970–6982 (2020).

40. Tong, Y. et al. Transcriptome-wide mapping of small-molecule rna-binding sites in cells informs an isoform-specific degrader of qsox1 mrna. J. Am. Chem. Soc. 144, 11620–11625 (2022).

41. Zhang, P. et al. Reprogramming of protein-targeted small-molecule medicines to rna by ribonuclease recruitment. J. Am. Chem. Soc. 143, 13044–13055 (2021).

42. Gupta, R. A. et al. Long non-coding rna hotair reprograms chromatin state to promote cancer metastasis. nature 464, 1071–1076 (2010).

43. Krishna, R. et al. Generalized biomolecular modeling and design with rosettafold all-atom. Science eadl2528 (2024).

44. Adasme, M. F. et al. Plip 2021: Expanding the scope of the protein–ligand interaction profiler to dna and rna. Nucleic acids research 49, W530–W534 (2021).

